# Genome-wide association mapping reveals genes underlying population-level metabolome diversity in a fungal crop pathogen

**DOI:** 10.1101/2022.05.20.492824

**Authors:** Nikhil Kumar Singh, Sabina Moser Tralamazza, Leen Nanchira Abraham, Gaétan Glauser, Daniel Croll

## Abstract

**Background:** Fungi produce a wide range of specialized metabolites (SMs) involved in biotic interactions. Pathways for the production of SMs are often encoded in clusters of tightly arranged genes identified as biosynthetic gene clusters. Such gene clusters can undergo horizontal gene transfers between species and rapid evolutionary change within species. The acquisition, rearrangement and deletion of gene clusters can generate significant metabolome diversity. However, the genetic basis underlying variation in SM production remains poorly understood.

**Results:** Here, we analyzed metabolite production of a large population of the fungal pathogen of wheat, *Zymoseptoria tritici*. The pathogen causes major yield losses and shows variation in gene clusters. We performed untargeted ultra-high performance liquid chromatography-high resolution mass spectrometry to profile the metabolite diversity among 102 isolates of the same species. We found substantial variation in the abundance of the detected metabolites among isolates. Integrating whole-genome sequencing data, we performed metabolite genome-wide association mapping to identify loci underlying variation in metabolite production (*i.e*. metabolite-GWAS). We found that significantly associated SNP reside mostly in coding and gene regulatory regions. Associated genes encode mainly transport and catalytic activities. The metabolite-GWAS identified also a polymorphism in the 3’UTR region of a virulence gene related to metabolite production and showing expression variation.

**Conclusions:** Taken together, our study provides a significant resource to unravel polymorphism underlying metabolome diversity within a species. Integrating metabolome screens should be feasible for a range of different plant pathogens and help prioritize molecular studies.

## Introduction

Fungi are capable of synthesizing a wide range of compounds including amino acids, pigments, antibiotics and toxins (Vining, 1990; Hoffmeister & Keller, 2007; Bouws *et al*., 2008). Such metabolites are typically classified as primary and specialized (or secondary) metabolites. Primary metabolites are essential for growth, development and reproduction and tend to be conserved across the phylogeny of fungi. Specialized metabolites (SM) confer benefits in specific ecological niches but are not essential for cell survival (Keller *et al*., 2005). The production of primary metabolites is encoded by a vast array of enzymes involved in metabolic reactions. In contrast, the production of SMs in fungi is often encoded by a small set of enzymes establishing a minimal metabolic pathway. The underlying genes tend to be arranged in close proximity forming a biosynthetic gene cluster (BCGs) including tight regulatory control (Keller, 2015). SMs play an important role in shaping species interactions with the microbiome. For example, the production of the antibacterial bikaverin is induced upon contact with the bacterium *Ralstonia solanacearum* is conserved in distant fungal pathogens (Spraker *et al*., 2018). Besides, SMs are essential for regulating fungal pathogenicity by acting as virulence factors against animal and plant hosts (Fox & Howlett, 2008). In the case of *Fusarium graminearum*, the BGC encoding the pathway for the production of the mycotoxin trichothecene is upregulated during colonization of wheat (Lysøe *et al*., 2011) and essential for fungal colonization of kernels. SM can also act as messengers in inter and intra-species communications between fungi (Yim *et al*., 2007; Netzker *et al*., 2015). The high degree of niche specificity of SMs generates diversity in BGCs among closely related species and even within some species (Keller, 2015, 2019). BGCs are also among the most mobile elements in fungal genomes with extensive evidence for horizontal gene transfer and turnover within species (Lind *et al*., 2015).

Large-scale fungal genome analyses have revolutionized the discovery of genes underlying the specialized metabolism and rapidly evolving BGCs transferred among species (Keller, 2019; Gil-Serna *et al*., 2020; Naranjo-Ortiz & Gabaldón, 2020). Genetic changes underlying the production of SM can be pinned down to single nucleotide variation as shown for fumonisin production in *Fusarium* fungi (Proctor *et al*., 2008). The evolution of BGCs often involves larger changes including the gain and loss of individual genes or entire clusters (Carbone *et al*., 2007; Tralamazza *et al*., 2019). BGCs are often located in more repetitive regions of the genome including subtelomeres (Palmer & Keller, 2010). Furthermore, BGCs in some groups of fungi are mainly regulated by epigenetic changes including chromatin modification (Collemare & Seidl, 2019). The fact that BGCs encode entire pathways for the production of SM, horizontal gene transfers effective means for species to acquire novel ecological functions (Cornell *et al*., 2007; Marcet-Houben & Gabaldón, 2010; Wisecaver *et al*., 2014). Horizontal gene transfers and rearrangements have led to substantial differences within and among closely related species in the content of BGCs and the potential to produce SMs. Among *Aspergillus* fungi, BGCs contribute to shared virulence profiles based on gliotoxin and fumitremorgin production and species-specific virulence *e*.*g*. due to fumagillin (Steenwyk *et al*., 2020). Genetic differences in qualitative and quantitative variation of SM production are hence likely under strong selection. Variation in metabolite production among conspecific isolates can be used for association mapping. Identifying associations between genetic variants in populations with relative levels of metabolite production is performed as metabolite genome-wide association studies (mGWAS) with a primary focus on the plant model *Arabidopsis thaliana* (Strauch *et al*., 2015). Recent applications on rice cultivars identified genes underlying flavonoid production (Peng *et al*., 2017) and contributions to the phenol-amides metabolic pathway (Dong *et al*., 2015). Metabolite variation was often mapped to a small number of major effect loci (Chen *et al*., 2014; Jacobowitz & Weng, 2020). The intra-specific variation in metabolite profiles, high-quality genomic resources and experimental tractability make fungi attractive models for genome-wide association mapping approaches.

The ascomycete *Zymoseptoria tritici* is a highly polymorphic fungal pathogen of wheat and shows a marked variability in BGCs among members of the species (Plissonneau *et al*., 2016, 2018; Badet *et al*., 2020). The pathogen causes yield losses of ∼ 5–30% depending on environmental conditions (Jørgensen *et al*., 2014; Fones & Gurr, 2015). Populations sampled across the world show evidence for the gain and loss of genes constituting BGCs (Hartmann & Croll, 2017). Comparisons of complete genome assemblies confirmed that BGCs are partly or entirely missing in some isolates (Badet *et al*., 2020) raising questions about the functional relevance of the encoded SMs. However, the role of SM in the lifecycle of *Z. tritici* remains poorly understood. A metabolome analysis of the wheat infection process showed that lipid metabolism is the initial energy source during leaf colonization prior to the induction of host cell death (Rudd *et al*., 2015). The pathogen upregulates BGCs mostly during the same transition to feeding from dead plant material *(i*.*e*. necrotrophic lifestyle) ∼10 days after infection (Rudd *et al*., 2015; Palma-Guerrero *et al*., 2016). Precursors of melanin were also found in the metabolite profile at this stage (Rudd *et al*., 2015). Melanin, which plays an important role in virulence, ultraviolet protection and anti-microbial resistance in fungi, is one of the best studied SM of *Z. tritici*. The locus underlying variation in melanin production was first identified using quantitative trait mapping and then confirmed to be a polyketide synthase gene cluster (Lendenmann *et al*., 2014; Krishnan *et al*., 2018). Among individual variation in melanin accumulation is governed by the insertion of a transposable element, which impacts the regulation of the BGC (Krishnan *et al*., 2018). Beyond melanin production, the metabolomic diversity within the species and the underlying genetic basis remains unknown.

Here, we take advantage of genome-wide association (GWA) mapping using the production of individual metabolites under standardized conditions as trait values. GWA mapping in *Z. tritici* has been used successfully to identify the genetic basis underlying virulence on different wheat cultivars, resistance to fungicides and temperature adaptation (Hartmann *et al*., 2017, 2020; Singh *et al*., 2021; Dutta *et al*., 2021). We focused on a single wheat field to establish a panel of 102 isolates showing considerable genetic diversity (Singh *et al*., 2020). We performed untargeted ultra-high performance liquid chromatography-high resolution mass spectrometry on individual fungal cultures growing under sterile conditions. We found considerable variation in metabolite profiles with the majority of metabolites showing abundance variation among isolates. GWA mapping revealed significantly associated loci in proximity to genes encoding functions related to transport and catalytic activity.

## Results

### Species-wide polymorphism in specialized biosynthetic gene clusters

A pangenome based on 19 completely assembled genomes of *Z. tritici* isolates collected from six continents and 13 different countries (Badet *et al*., 2020) was used to retrieve between 29-33 BGCs per genome (Figure 1A). Non-ribosomal peptide synthetase (NRPS) and type 1-polyketide synthase are the most abundant gene clusters. Approximately 72% of the predicted core biosynthetic genes are conserved among isolates and ∼24% are accessory (present in 2-18 isolates) (Figure 1B). We also retrieved a singleton core biosynthetic gene found only in an isolate sampled in Tunisia. We found similar proportions for additional biosynthetic genes with ∼60% and 30% core and accessory genes, respectively (Figure 1B). Regulatory genes of gene clusters were all conserved, but only 90% of the transporter genes were conserved (Figure 1B). BGCs show polymorphism also at the level of single fields. Focusing on BGCs encoded in the genomes of four isolates collected in the same year from nearby wheat field in central Europe, seven clusters showed presence/absence variation (Badet *et al*., 2020) (Figure 1C).

**Figure 1:**
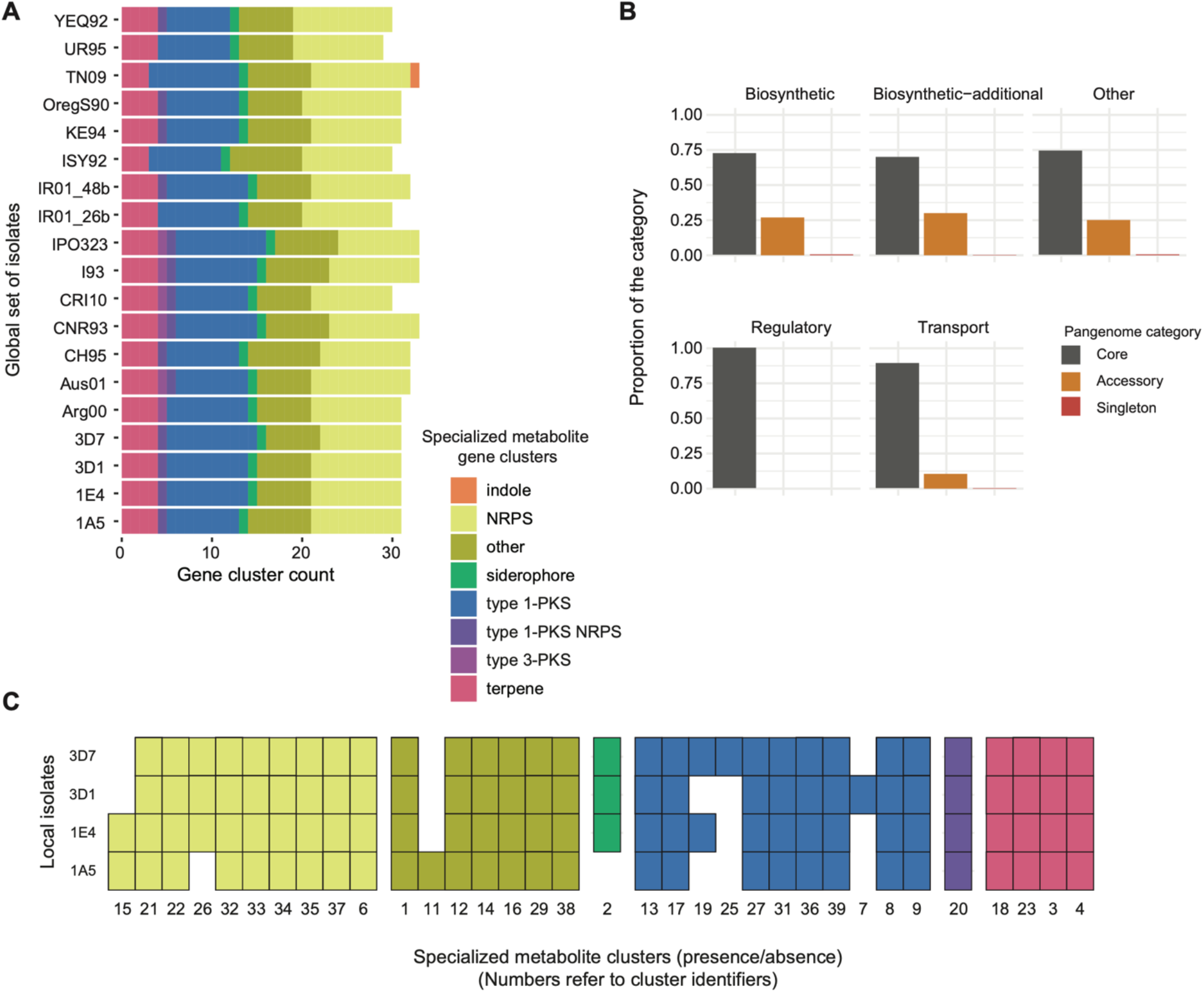
Genetic diversity of specialized metabolite gene clusters in *Zymoseptoria tritici*. A) Detected gene clusters in 19 genomes constituting the species pangenome (Badet *et al*., 2020). Colors indicate different categories of specialized metabolite gene clusters depending on the metabolite compound or the core biosynthetic gene: indoles, non-ribosomal peptide synthetase (NRPS), siderophores, polyketide synthase (PKS) and terpenes. B) The proportions of core, accessory and singletons specialized metabolite gene clusters in the species pangenome. C) Presence-absence variation of 39 gene clusters in genomes of four isolates collected in Switzerland.

### Genetic diversity in a single-field mapping population

To identify whether individual isolates differ in the production of metabolites and show heritable genetic variation, we performed a genome-wide metabolite association study (mGWAS). We analyzed whole genome sequences of 102 strains isolated from a single wheat field. The mapping population was previously shown to be well-suited for GWAS (Singh *et al*., 2021). The isolates were collected from 11 genetically different winter wheat cultivars (1-9 isolates per cultivar) from three collection time points (Figure 2A, Supplementary Table S1). At the first collection time point, the wheat seedling was at growth stage (GS) 41 where flag leaves start extending. At the second (GS 75) and third collection (GS 85) time point, plants were fully developed and grains were reaching maturity (Meier, 2001). The average Illumina sequencing coverage was 21X as previously described (Singh *et al*., 2020). After quality filtering, we retained 504,557 high-confidence single nucleotide polymorphisms (SNPs). We constructed an unrooted phylogenetic network using Splitstree and we found isolates to be at similar genetic distances (Figure 2B). The mapping population also included eight clonal groups for a total of 19 isolates. We used relatedness as a random factor in association mapping to account for the uneven relatedness. A principal component analysis (Figure 2C) identified a small group of outlier isolates. No genetic structure was found among collection time points or cultivars (Singh *et al*., 2020).

**Figure 2:**
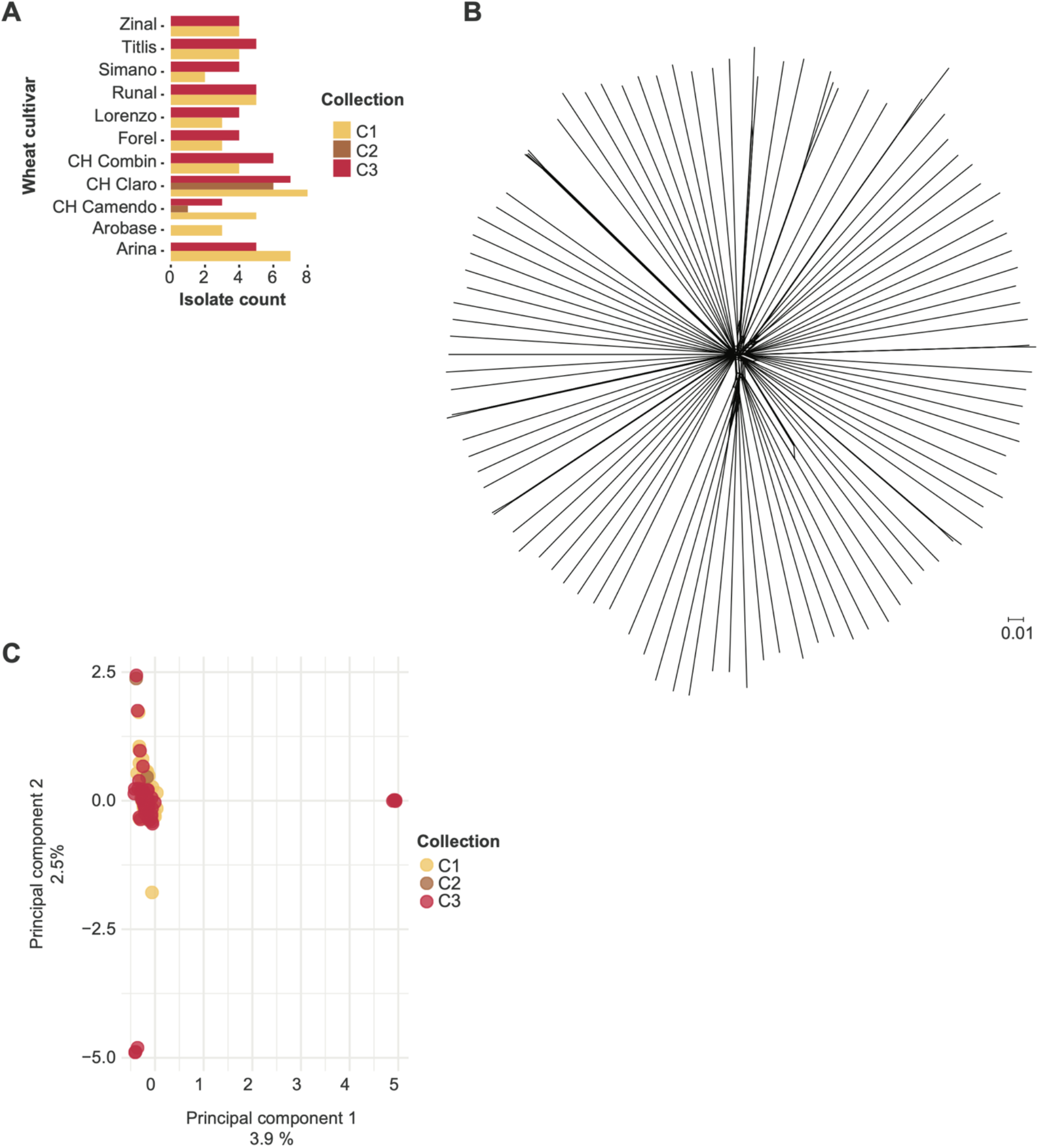
Whole-genome sequence analyses of 102 *Zymoseptoria tritici* isolates collected from a single field. A) Number of isolates collected from each of the eleven cultivars at each collection time point (C1-3; early, middle and late in the season; see also (Singh *et al*., 2020). B) A SplitsTree phylogenetic network constructed from genome-wide single nucleotide polymorphism (SNP) data. C) The first two principal components (PC) from a PC analysis of genome-wide SNPs. Isolates are color coded by the collection time point.

### Untargeted metabolite profiling at the population level

We performed an untargeted metabolite profiling of all 102 isolates using Ultra-high-performance liquid chromatography high-resolution mass spectrometry (UHPLC-HRMS). The analyzed metabolites were extracted from blastospores after culture washing. After excluding highly hydrophilic (*e*.*g*. sugars, amino acids) and hydrophobic (*e*.*g*. lipids) molecules based on retention times, we retained 2633 metabolite marker peaks (Supplementary Table S2). Using relative abundance across all peaks, we performed a principal component analysis (Figure 3A). Interestingly, the metabolite profiles of different isolates clustered based on their field collection time point (Figure 3A). This contrasts with the genome-wide differentiation at neutral markers (Figure 2C). We performed a down sampling analysis to identify the proportion of metabolite markers detected in isolates. We found approximately equal proportions of metabolite markers detected in 75%, 50% and 25% of the total isolates (Figure 2C).

**Figure 3:**
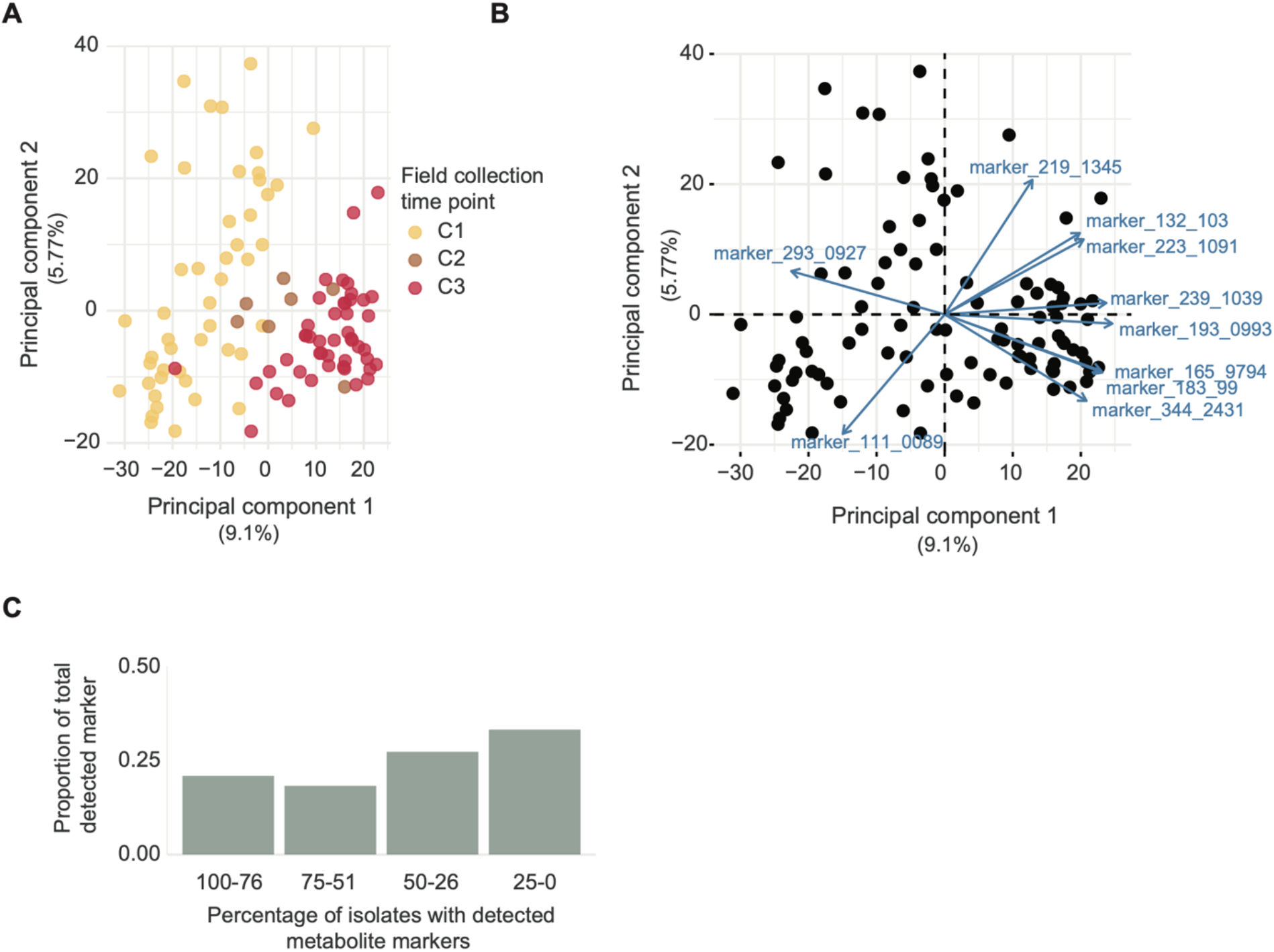
Untargeted metabolite profiling of 102 isolates using ultra-high performance liquid chromatography tandem mass chromatography (UHPLC-QTOF-MS). A) The first two principal components (PC) from a PC analysis of 2633 metabolite markers. Isolates are colour coded by the collection time point. B) Bi-plot of 2633 metabolite markers showing the contribution of the ten metabolite markers contribution most to differentiation of metabolite profiles. Metabolite markers are labelled using their respective *m/z* values. C) Down sampling analysis to identify the proportion of 2633 metabolite markers detected in mapping population

### Metabolite-GWAS based on variation in metabolite abundance

Variation in relative abundance among isolates may have a genetic basis (Figure 3A). We performed mGWAS on relative metabolite marker intensities based on mixed linear models taking genetic relatedness into account. We first evaluated top association *p*-values for 2388 untargeted metabolite markers showing distinct *m/z* peaks. After filtering association *p*-values, 68.8% (1644 out of 2388) of untargeted metabolites showed at least one significant SNP association (Supplementary Table S3). Overall, we found similar proportions of associated SNP on core and accessory chromosomes (Figure 4A). Interestingly, accessory chromosome 14 showed a higher proportion of associated SNPs. Chromosome 14 carries a substantial insertion in some isolates of the species including coding sequences (Croll et al. 2013, Badet et al 2020).

**Figure 4:**
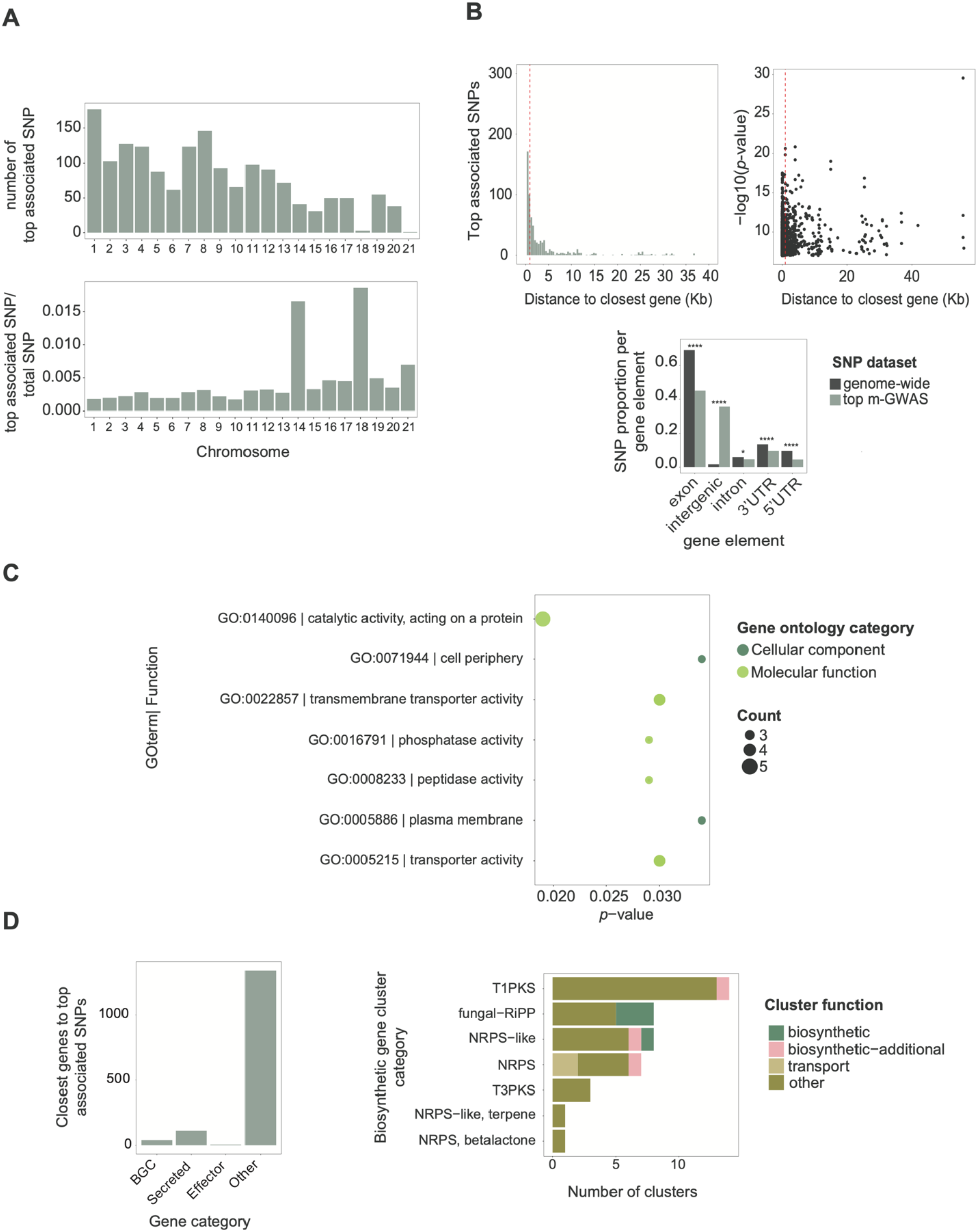
Metabolite genome-wide association studies (mGWAS) and candidate gene functions. A) Number of the most significantly associated single nucleotide polymorphisms (SNP) and proportion of significantly associated SNPs. B) Distance to the closest gene and association −log10(*p*) values of the most significantly associated SNP for each metabolite marker. The dotted red line delimits 75% of the SNPs. Proportion of SNPs found per gene element across the genome and for the significantly SNPs. C) Gene ontology term enrichment analysis of functions encoded by the closest genes. D) Number of the most significantly associated SNP to the closest gene based on gene type, biosynthetic gene cluster class and function. *, **** : *p-*value <0.05, <0.00001, respectively.

We analyzed the closest gene to each significantly associated SNP (Figure 4B, Supplementary Table S3). Overall, 75% of the most significantly associated SNP for each metabolite marker fell was found at a maximum distance of 915 bp to the closest gene and 50% within 18 bp (Figure 5B). The significantly associated SNPs were highly enriched for intergenic regions (*p <* 0.00001) indicating that such variants may play a regulatory role. Linkage disequilibrium on chromosomes decays to *r*^2^ < 0.2 within 500-1500 bp (Singh *et al*., 2020). This range matches also the average gene distance in the *Z. tritici* genome (Goodwin *et al*., 2011). For the remainder, we focused only on the most significantly associated SNPs for each metabolite marker falling within 5 kb of a gene (Supplementary Table S3, *n* = 1508).

**Figure 5.**
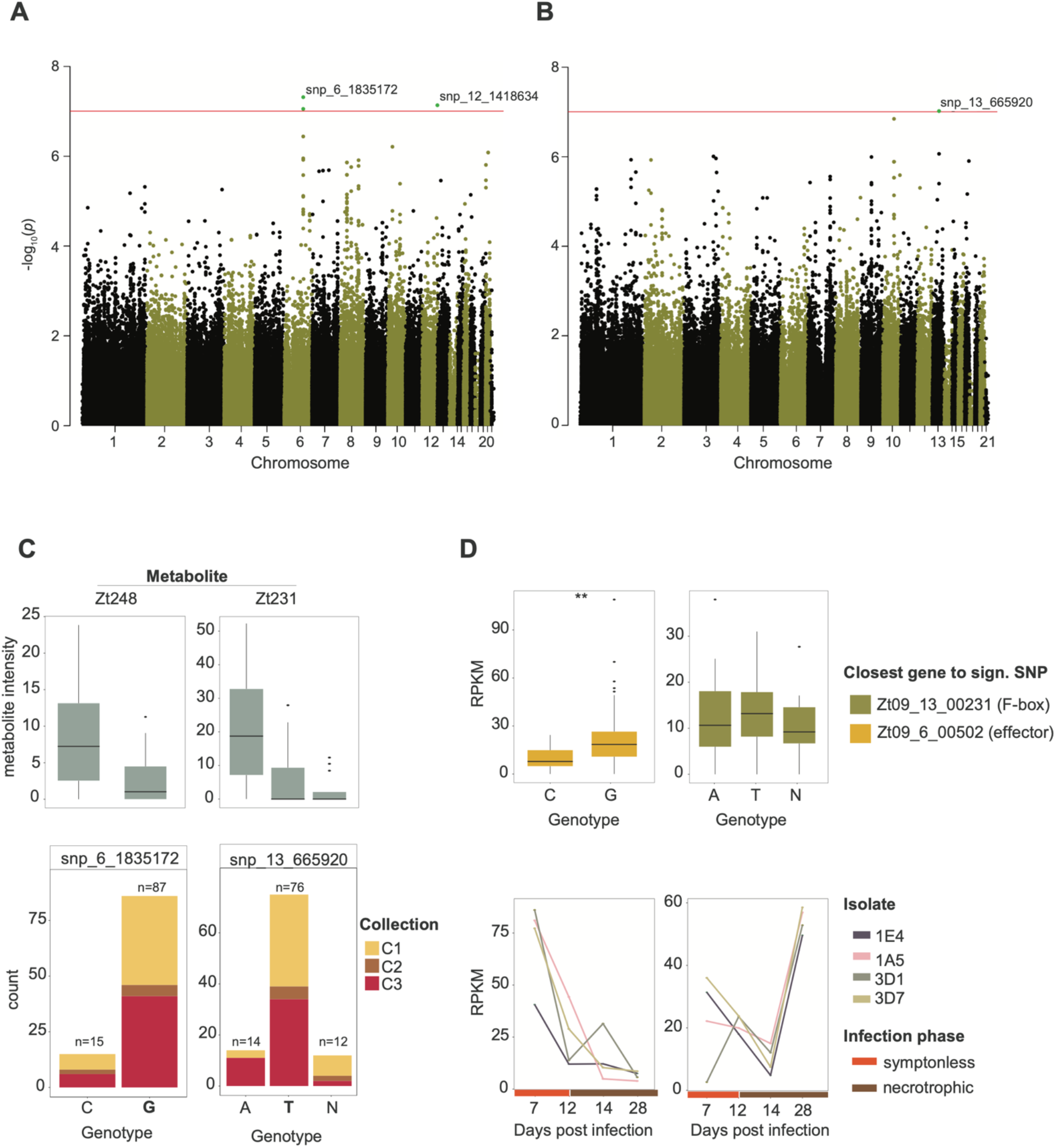
Analyses of key loci identified by the metabolome-GWAS. A) Manhattan plots of genome-wide association mapping performed for the metabolite Zt248 and (B) the metabolite Zt231 in a *Z. tritici* single-field population (*n*=102 isolates). The red line refers to the Bonferroni threshold (α < 0.05). C) Relative metabolite intensity and isolate counts based on genotypes (N: unassigned genotype). Colors refers to field collection time point of the isolates. D) Expression of the closest genes to the focal SNP under culture conditions (same isolates as mapping population; upper box) and of four different isolates collected nearby over the course of an experimental wheat infection (lower box).

We performed gene ontology (GO) term enrichment analyses of the protein functions encoded by genes nearest to the most significantly associated SNP (Supplementary Table S4). We found that 27% of all nearby genes were assigned GO terms in contrast with 49% for all genes in the genome (Supplementary Table S3). We found the strongest enrichment for membrane transport functions and catalytic activity (Figure 4C, Supplementary Table S4). Transporters play key roles in metabolic pathways (Keller, 2019) and often underpin niche adaptation (Stewart et al. 2018).

### Metabolite-GWAS candidates targets

We focused on the most significantly associated SNPs closest to genes. We found 124 genes encoding predicted proteins including seven proteins characterized as putative effectors (Figure 4D). We additionally found 42 genes involved in specialized biosynthetic gene clusters (BGC) targeting 19 out of the 39 known BGC in the species (Figure 1A). BGCs of the polyketide, fungal-RiPP, and NRPS classes showed the most associations (Figure 4D). Next, we manually curated promising targets based on metabolite peak quality, relative marker intensity variation, and SNPs to gene distance for a maximum of 500 bp away from coding sequences.

The biallelic SNP at 1,835,172 bp on chromosome 6 showed a significant association with the metabolite Zt248 (m/z of 248.1323 and retention time of 3.52 min; Figure 5A). We found that 14% of the population carries the alternative C allele associated with higher metabolite production (Figure 5B). We found no association of the C allele with the field collection time point (Figure 5C). The SNP is 180 bp downstream (3’UTR) of a functionally predicted effector gene Zt09_6_00502 (Supplementary Table S3). Based on RNA-seq analyses of the isolates growing in liquid culture, we found that a significant association between transcription of gene Zt09_6_00502 and the genotypes at SNP discovered in the mGWAS (*p-value* < 0.001; Figure 5D). Hence, the locus may mediate metabolite production through variation in gene expression. Furthermore, the predicted effector is primarily expressed during the initial phase of the wheat infection (Figure 5D) suggesting contributions to the onset of the disease. Interestingly, Zt09_6_00592 is identified as a component of a NRPS gene cluster (cluster 18; Figure 1A). The gene cluster is 35kb in size and is predicted to carry 15 genes including two biosynthetic core genes and a regulatory gene (Figure 6A). The gene cluster is conserved within the *Z. tritici* and shared between three closely related sister species (Figure 6A).

**Figure 6.**
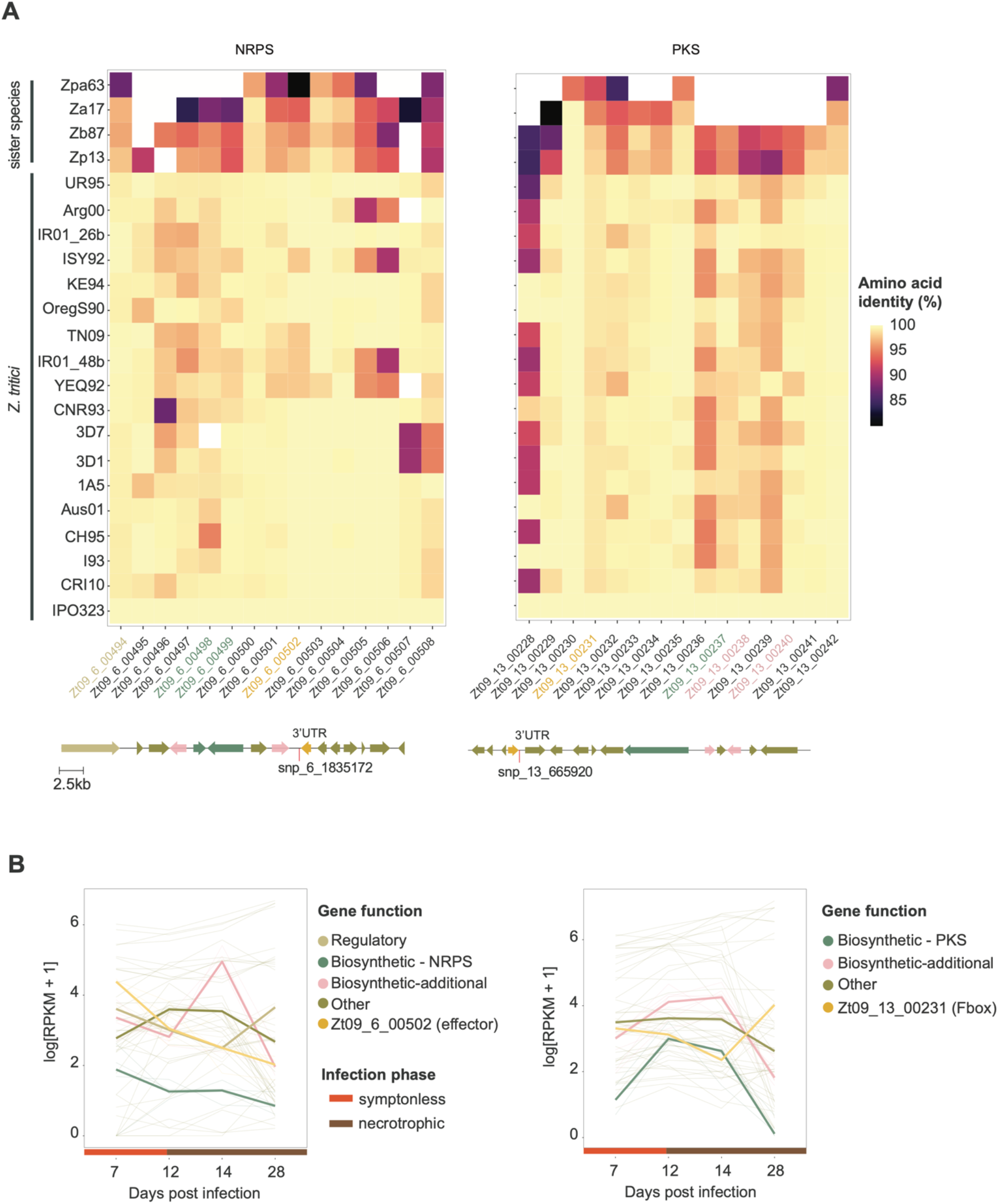
Conservation of the NRPS and PKS biosynthetic gene clusters in the *Z. tritici* pangenome and sister species. A) Conservation of the effector Zt_06_00502 gene in the NRPS gene cluster and the F-box gene Zt_13_00231 in the PKS gene cluster. Amino acid identity compared to the gene cluster in the *Z. tritici* reference genome IPO323. Arrows below the heatmap correspond to genes locations and SNP positions in the reference genome. B) Mean expression of the NRPS and PKS gene clusters based on four isolates collected in a nearby field (3D1, 3D7, 1A5 and 1E4). Colors identify gene functions. NRPS: nonribosomal peptide-synthetase, PKS: polyketide synthase. Sister species isolates Zp13: *Z. pseudotritici*, Zpa63: *Z. passerinii*, Zb87: *Z. brevis*, Za17: *Z. ardabiliae*.

We found that the biosynthetic core genes share a similar transcription profile as the effector gene indicating co-regulation during the infection cycle. Furthermore, we analyzed data on controlled infections based on the same set of isolates and found that the isolates carrying the alternative genotype caused consistently more lesions (median of 71% of leaf area covered by lesion) in contrast to strains carrying the reference genotype (median of 57 % of leaf area covered by lesions; Supplementary Figure S1) consistent with higher metabolite production contributing to host damage. We performed a multiple sequence alignment of the 3’UTR region containing the significantly associated SNPs using species across the *Zymoseptoria* genus. The phylogenetic context of the 3’UTR indicates that the alternative allele is derived in *Z. tritici*. Interestingly, variants in the gene sequence of Zt09_6_00502 are consistent with geographic differentiation of the species. In contrast, variants in the 3’UTR region reveal haplotypes shared between the Swiss field population (ST16CH_1A27 and ST16CH_1M28) and geographically distant isolates from Yemen (Yeq92) and Tunisia (TN09; Supplementary Figure S2).

We investigated a second locus in detail located at 665,920 bp on chromosome 13 with a significant association with metabolite Zt231 (m/z 231.171 and retention time of 2.24 min; Figure 5B). Isolates carrying the alternative allele A showed higher metabolic intensity relative to isolates carrying the reference allele T (Figure 5C). The SNP is located 255 bp downstream (3’UTR) of the F-box gene Zt09_13_900231. The encoded F-box domain (position 55-99) overlaps with a leucine rich region (LRR domain; amino acid position 42-374; Supplementary Figure S3). Expression analyses of Zt09_13_900231 revealed no significant association with the genotypes identified through mGWAS under culture conditions (Figure 5D). However, we identified an expression quantitative trait locus (eQTL) mapped using the same single-field population at the same mGWAS locus (Abraham et al.; snp_13_664471). The transcription of the F-box Zt09_13_900231 gene was upregulated at the begging of the infection (7 dpi) followed by a decrease during the transition to the necrotrophic phase (12-14 dpi) and a final increase at 28 dpi late in the infection (Figure 5D). The gene Zt09_13_00231 is predicted to encode a component of the PKS gene cluster 18 (Figure 1B). The cluster is conserved within *Z. tritici* and the sister species *Z. brevis* (Figure 6A). The gene cluster includes a PKS core gene and two additional biosynthetic genes. Interestingly, we found that the transcription of the BGC is largely antagonistic to the transcription of the F-box gene (Zt09_00231) suggesting negative feedback mechanism during infection (Figure 6B). Experimental infections of wheat showed a trend of higher pycnidia production of the isolates carrying the alternative genotype associated with higher metabolite production (Supplementary Figure S1).

### Chemical characterization of associated metabolites

We analyzed the two metabolites (Zt248 and Zt231) further using chemical and natural product databases. The prediction based on the software Canopus (Dührkop et al. 2020) classified the metabolite Zt248 (C_11_H_21_NO_3_S) with 90% confidence as a cysteine derivative. The Pubchem database classified Zt248 as an N-Acetyl-S-(2-ethylbutyl)-L-cysteine. The metabolite Zt231 (C_11_H_22_N_2_O_3_) produced fragments at m/z 86.097 typical for leucine and isoleucine moieties. A natural product database search retrieved N-Valyl-Leucine as the most likely compound for Zt231. Our analysis using MS/MS showed three peaks at 2.15, 2.24 and 2.37, corresponding to the D-l, L-D and L-L isoforms (Supplementary Figure S4). We searched for possible analogues and found that confluenine A produced by the basidiomycete *Albatrellus confluens* (Zhang et al. 2018) shares the same molecular formula. The underlying polyketide BGC shares no homology among known fungi though. According to a CFM-ID *in-silico* fragmentation confluenine A should fragment into m/z 85.065 and metabolite Zt231 should fragment to m/z 85.02 and 86.09.

## Discussion

We used untargeted metabolite profiling and genome-wide association mapping to identify candidate loci underlying the production of metabolites. We found that a single, highly diverse field population of *Z. tritici* likely harbors substantial variation in the production of individual compounds. We performed association mapping based on standardized metabolite peak intensities as a phenotypic trait and identified significantly associated loci in close proximity to regulatory regions encoding virulence factors and contain biosynthetic gene clusters.

We identified highly variable metabolite profiles among isolates from a single field population. Given the standardized conditions, genetic factors are likely contributing to the observed metabolome diversity. The analyzed population is genetically highly diverse with only weak genetic substructure (Singh *et al*., 2020). Surprisingly, overall metabolite profiles clustered according to the isolate collection time point over the growing season (*i*.*e*. C1-C3 spread over several weeks). This is in contrast to the genetic differentiation of the same isolates where no clustering is evident based on collection (Singh *et al*., 2020). A possible explanation for the consistent shifts of the metabolome over the course of the growing season is selection for isolates expressing a specific set of metabolites. A possible environmental factor driving selection are the two fungicide treatments applied midway between the collections C1 and C2 (Singh et al. 2021). Additional shifts in phenotypic traits were also apparent for pathogen reproduction (*i*.*e*. pycnidia formation) on wheat leaves with isolates collected later in the season showing higher reproductive output under controlled conditions (Singh *et al*., 2020, 2021). Shifts in aggressiveness were also previously observed in multi-year studies of *Z. tritici* over years, cultivars and leaves on individual plants (Suffert *et al*., 2018). Identifying the chemical structure of the compounds contributing most strongly to shifts in the metabolome over the course of the growing season will provide insights into the adaptive nature of such changes.

Variation in metabolite abundance among individuals can be used to identify candidate genes underlying the regulation of metabolite production. We found a strong enrichment of intergenic regions contributing to metabolite production suggesting *cis*-regulation (*i*.*e*., promoter regions) playing a role in shaping within species metabolome diversity. We identified genetic polymorphisms close to genes encoding a broad range of functions with an enrichment of transporter and catalytic functions. Such transporters include major facilitator superfamily (MFS) transporters known to modulate fungicide and stress tolerance (Roohparvar *et al*., 2007; Omrane *et al*., 2017; Zhong *et al*., 2021). MFS transporters are also involved in the secretion of phytotoxins during infection and, hence, can contribute to virulence (Choquer *et al*., 2007; Temme *et al*., 2012; Menke *et al*., 2012).

The study species harbors a high degree of polymorphism in BGCs with likely consequences for the production of specialized metabolites. The mGWAS captured 41 BGC-related associations affecting approximately half of all known BGCs of the species. Standing variation for the production of SM elevates the evolutionary potential of *Z. tritici*, because populations could respond rapidly to selection for or against the production of specific metabolites. Intra-species variation in BGCs and SM production have been observed in a range of fungi including the rice blast pathogen *Magnaporthe oryzae*, where a gain of virulence was linked to the duplication of a hybrid PKS-NRPS cluster (Zhong *et al*., 2020). *Aspergillus* species show also significant intra- and interspecific variation in BGCs and the potential to produce specific SM with consequences for niche adaptation (Theobald *et al*., 2018).

The *in vitro* standardized mGWAS successfully captured targets associated with host infection. The candidate effector gene Zt09_00502 belongs to the NRPS gene cluster 18 upregulated during the initial phase of infection. Pathogens secrete effectors to manipulate the host immune responses and physiology to its advantage (Sánchez-Vallet, et al., 2018). The production of the metabolite is also associated with the expression of the candidate effector. Infection data indicated higher aggressiveness of the isolates producing higher levels of the metabolite. The conservation of the gene cluster including the effector suggestions though that the cluster function has gained its original function prior to the specialization of the pathogen on wheat. However, the recently arisen adaptive mutation associated with the metabolite production could be an evolutionary response of the pathogen to gain an advantage on wheat.

The second investigated metabolite association was near the gene Zt09_13_00231 encoding an F-box protein, which generally bind and transport proteins to be discarded by the cell (Patton et al., 1998). The encoded protein contains domains consistent with a typical F-box FBLX family protein shown to be involved in conidiation, carbon intake and virulence regulation in fungi (Han et al., 2007, Guo et al., 2015). Deletion of the gene encoding a homologous F-box MoGrr1 in the fungal pathogen *Magnaporthe oryzae* resulted in reduced lesion sizes on rice (Shi et al., 2019). Chemical structure predictions and database searches revealed that the metabolite Zt231 associated with the F-box gene shares a structure matching confluenine A. This metabolite has antimicrobial activities and is produced by the basidiomycete *A. confluens* (Zhang et al., 2018). Convergent molecular structures may be an indicator of similar functionality, however detailed chemical analyses and interaction studies on the plant are needed.

In conclusion, our study highlights the power of association mapping to identify candidate loci underlying metabolite diversity and associated with the host infection process. Previously, mGWAS was limited to plant models, but our findings illustrate how applications to plant pathogenic fungi can efficiently produce candidate lists. A key requirement is access to large, experimentally tractable isolate collections. High degrees of recombination and rapid decay of linkage disequilibrium are further requirements for the successful application of association mapping approaches (Sánchez-Vallet *et al*., 2018). Establishing metabolome-wide profiles for a range of pathogens under variable conditions will complement analyses focusing on proteaceous effectors.

## Methods

### Field collection and storage

We collected *Z. tritici* isolates from the Field Phenotyping Platform (FIP) site of the ETH Zürich, Switzerland (Eschikon, coordinates 47.449°N, 8.682°E) (Kirchgessner *et al*., 2017). We analyzed a total of 102 isolates collected during the 2016 growing season from 11 winter wheat cultivars, which are commonly grown in Switzerland (Levy et al., 2017). We analyzed isolates originating from three collection time points over the season (Supplementary Table S1). The first collection (*n* = 48) was established in May when wheat plants were in growth stage (GS) 41. The second (*n* = 7) and third collection (*n* = 47) was established when the plants were in GS 75 and GS 85, respectively. After sampling, spores of each isolate were stored in either 50% glycerol or anhydrous silica gel at −80 °C. Additional information regarding the sampling scheme and genetic diversity of the collection is described in Singh *et al*. (2020)

### Culture preparation and metabolite extraction

Isolates were revived from glycerol stock by adding 50 μl fungal stock solution to a 50 ml conical flask containing 35 ml liquid YSB (yeast-sucrose broth) medium. The inoculated flasks were incubated in the dark at 18° C and 160 rpm on a shaker-incubator. After 8 days of incubation, the cultures were passed through four layers of meshed cheesecloth and washed thrice with sterile milli-Q water to remove media traces. Spores were then lyophilized and metabolites extracted by resuspending the spores (∼80 mg) in 1 ml of HPLC-grade methanol. The extract was centrifuged at 15,000 rpm for 5 minutes to pellet down debris before retrieving the supernatant. The last step was repeated until a clear supernatant was recovered.

### Whole-genome sequencing and variant calling

Approximately 100 mg of lyophilized spores were used to extract high-quality genomic DNA with the Qiagen DNeasy Plant Mini Kit according to the manufacturer’s protocol. We sequenced paired-end reads of 100 bp each with an insert size of ∼550 bp on the Illumina HiSeq 4000 platform. Raw reads are available on the NCBI Sequence Read Archive under the BioProject PRJNA596434 (Oggenfuss *et al*., 2020). Illumina sequences were quality checksed using FastQC v. 0.11.9. (Andrews, 2010). Sequencing reads were then screened for adapter sequences and quality trimmed using Trimmomatic v. 0.39 (Bolger et al., 2014) using the following settings: illuminaclip=TruSeq3-PE.fa:2:30:10, leading=10, trailing=10, sliding-window=5:10 and minlen=50. Trimmed sequencing reads were aligned to the reference genome IPO323 (Goodwin et al., 2011) available from Ensembl Fungi (https://fungi.ensembl.org/Zymoseptoria_tritici/Info/Index) and the mitochondrial genome (European Nucleotide Archive accession EU090238.1) using Bowtie2 v. 2.4.1 (Langmead & Salzberg, 2012). Multi-sample joint variant calling was performed using the HaplotypeCaller and GenotypeGVCF tools of the GATK package v. 4.0.1.2 (McKenna et al., 2010). We retained only SNP variants (excluding indels) and proceeded to hard filtering using the GATK VariantFiltration tool based on the following cutoffs: QD < 5.0; QUAL < 1000.0; MQ < 20.0; -2 > ReadPosRankSum > 2.0; -2 > MQRankSum > 2.0; -2 > BaseQRankSum > 2.0. After filtering for locus level genotyping rate (>50%) and minor allele count (MAC) of 1 using VCFtools v. 0.1.15 (Danecek et al., 2011). Additional sequencing and variant call statistics are available from Singh *et al*. (2021).

### Untargeted metabolite profiling using UPLC-HRMS

Metabolome analyses were carried out by UHPLC-HRMS using an Acquity UPLC coupled to a Synapt G2 QTOF mass spectrometer (Waters). An Acquity UPLC HSS T3 column (100×2.1mm, 1.8 μm; Waters) was employed at a flow rate of 500 μl/min and maintained at a temperature of 40°C. The following gradient with 0.05% formic acid in water as mobile phase A and 0.05% formic acid in acetonitrile as mobile phase B was applied: 0-100 % B in 10 min, holding at 100% B for 2.0 min, re-equilibration at 0% B for 3.0 min. The injection volume was 2.5 μl. The QTOF was operated in electrospray negative mode using data-independent acquisition (DIA) alternating between two acquisitions functions, one at low and another at high fragmentation energies. Mass spectrometric parameters were as follows: mass range 50-1200 Da, scan time 0.2 s, source temperature 120°C, capillary voltage -2.5 kV, cone voltage -25V, desolvation gas flow and temperature 900 L/h and 400°C, respectively, cone gas flow 20 L/h, collision energy 4 eV (low energy acquisition function) or 15-50 eV (high energy acquisition function). A 500 ng/ml solution of the synthetic peptide leucine-enkephaline in water:acetonitrile:formic acid (50:50:0.1) was infused constantly into the mass spectrometer as internal reference to ensure accurate mass measurements (<2ppm). Data was recorded by Masslynx v.4.1. Marker detection was performed using Markerlynx XS (Waters) with the following parameters: initial and final retention time 1.5 and 10.0 min, mass range 85-1200 Da, mass window 0.02 Da, retention time window 0.08 min, intensity threshold 500 counts, automatic peak width and peak-to-peak baseline noise calculation, deisotoping applied. Data was mean-centered and Pareto scaled before applying multivariate analysis. Markers of interest were characterized using the mass spectrometry reference spectra database MassBank (https://massbank.eu).

#### Genome-wide association mapping and linkage disequilibrium analyses

We performed GWAS based on mixed linear models accounting for degrees of kinship among genotypes (MLM+K). The kinship matrix was computed using the scaled identity-by-state (IBS) algorithm implemented in TASSEL v. 20201114 (Bradbury *et al*., 2007). We included the kinship matrix as a random effect in the mixed linear models for association mapping using TASSEL. Accounting for kinship performs sufficiently well to control for genetic substructure in the mapping population (Singh *et al*. 2021). Untransformed relative abundance values for each peak were used as trait values for association mapping. Outcomes were visualized using the R package *qqman* v. 0.1.4 (Turner, 2014). We filtered association *p*-values based on the Bonferroni threshold at α = 0.05. The closest gene for each associated SNP was identified using the “closest” command in bedtools v. 2.29.0 (Quinlan & Hall, 2010). GO term enrichment analyses were performed using the Fisher’s exact test based on gene counts with the topGO R package (Alexa & Rahnenfuhrer, 2009) and plotted using the GOplot R package(Walter et al., 2015).

## Acknowledgements

We would like to thank Ophélie Gning for providing technical support in the metabolite sample preparation. The research was funded by a Swiss National Science Foundation grant to DC (number 173265 and 177052).

## Notes

### Competing Interest Statement

The authors have declared no competing interest.

