## Supplementary Figure for "Genome-wide association mapping reveals genes underlying population-level metabolome diversity in a fungal crop pathogen"

**Supplementary Figures**

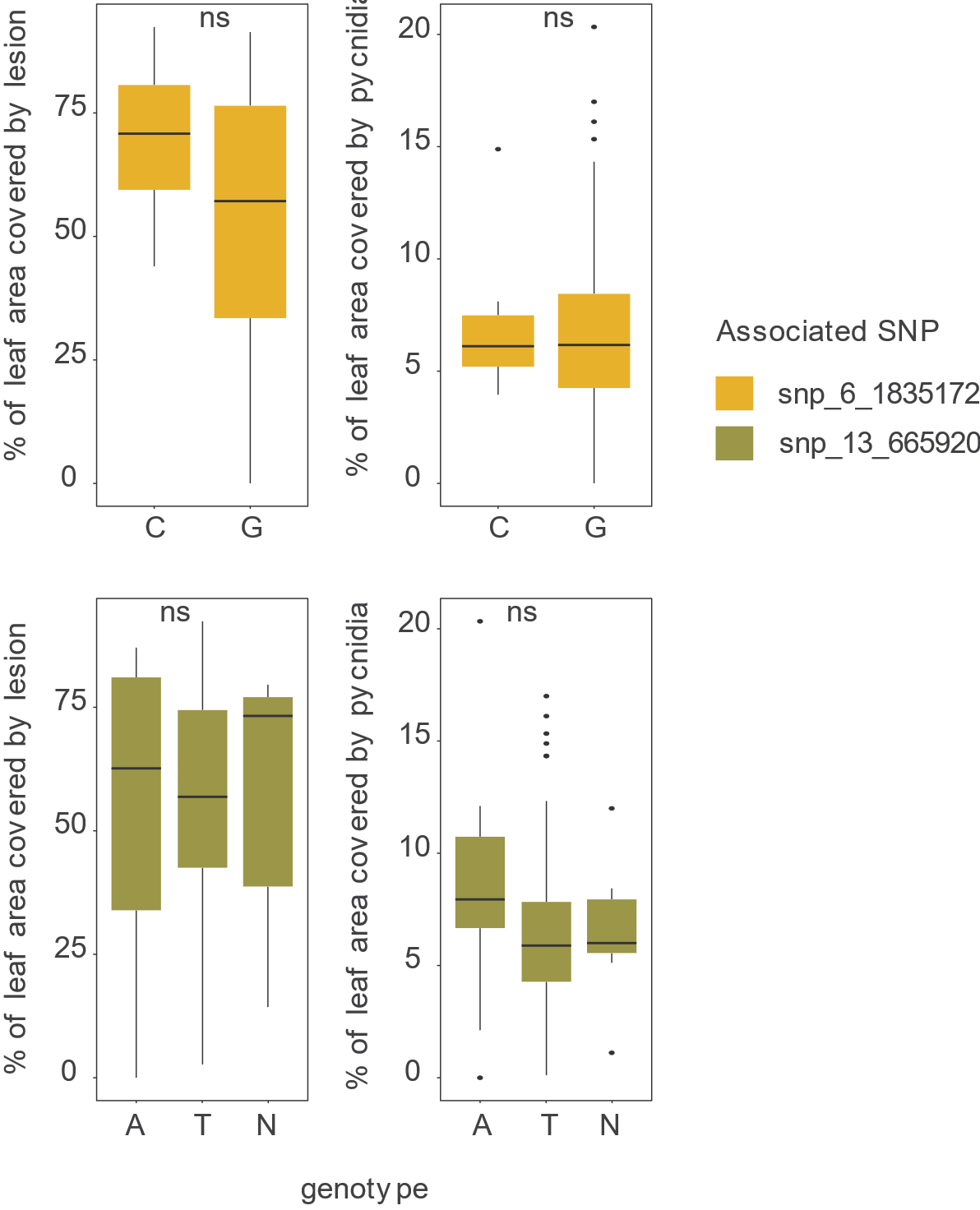

**Supplementary Figure S1**. Percentage leaf area covered by lesions and pycnidia during wheat infection for *Z. tritici* strains included in the metabolome-GWAS analysis (*n* = 102). The isolates are grouped by their genotype at significant metabolome GWAS SNPs.

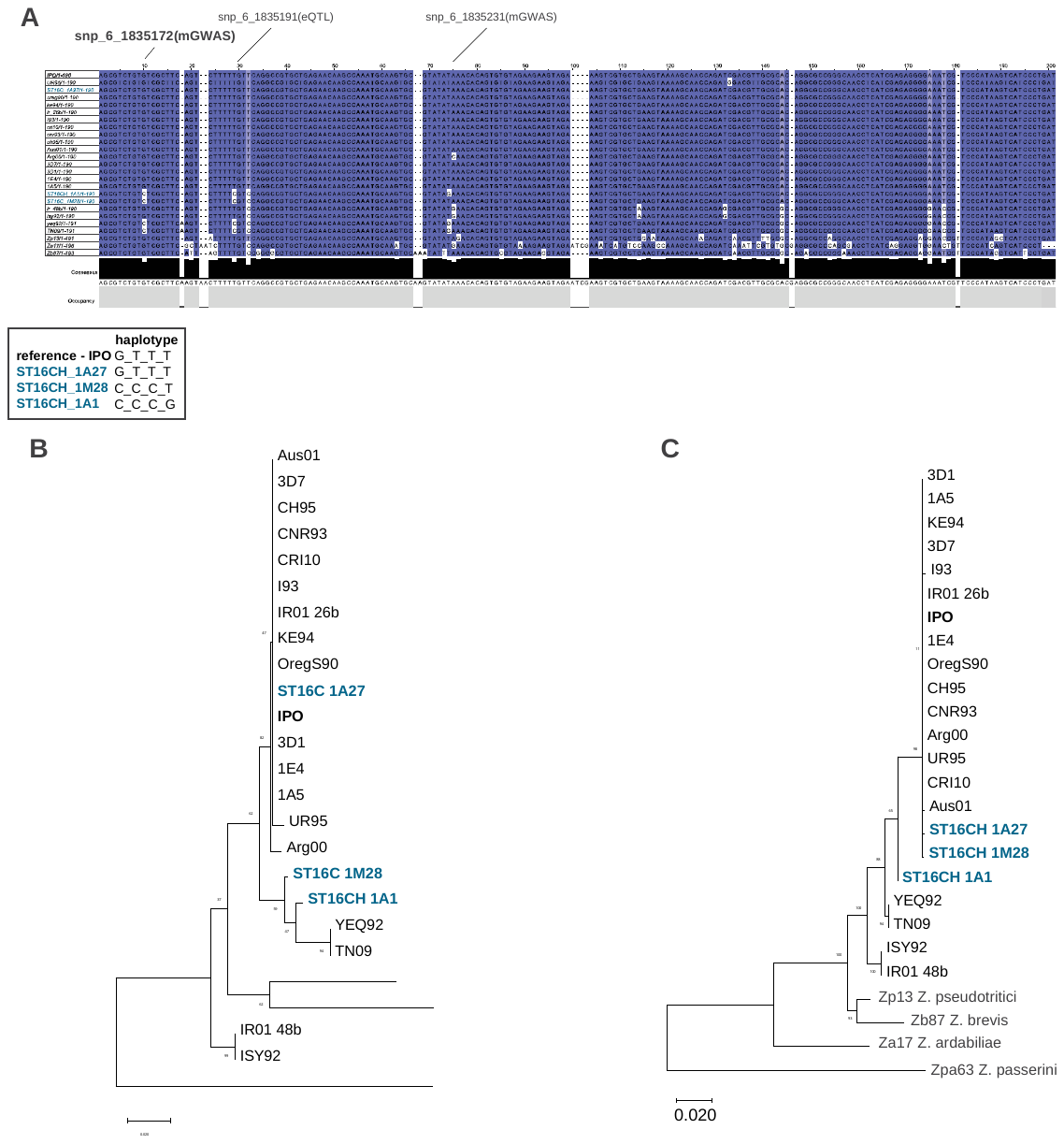

**Supplementary Figure S2.** Evolutionary history of the putative effector gene *Zt09_00502*. A) Alignment of the 3’UTR region of the gene. The box refers to the haplotypes found in the single Swiss field population. B) Phylogenetic tree of the 3’UTR region and (C) of the gene sequence. Names in bold black, blue and grey refer to the reference genomes of the species (IPO), isolates included in the metabolite GWAS and sister species, respectively. The phylogenetic tree was inferred by using maximum likelihood and the Tamura-Nei model.

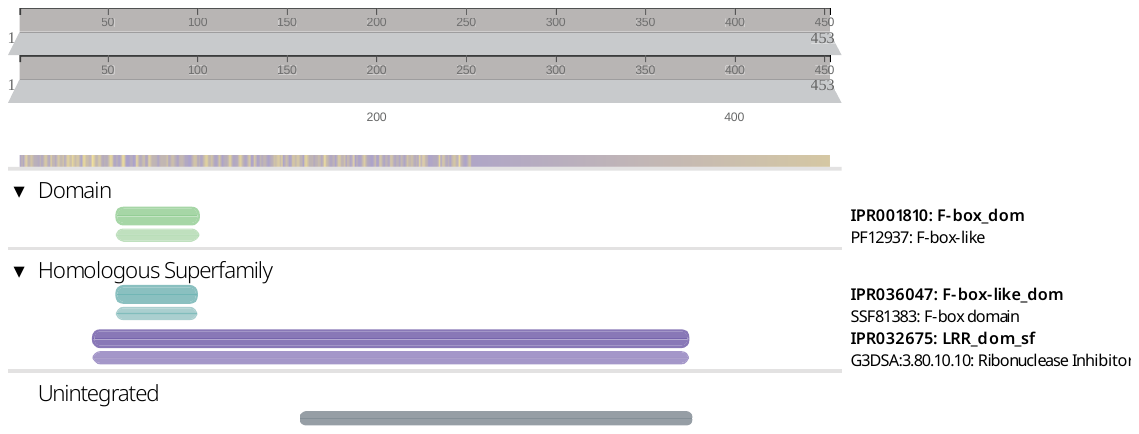

**Supplementary Figure S3.** Classification of protein family domains encoded by the gene *Zt09_13_00231.*

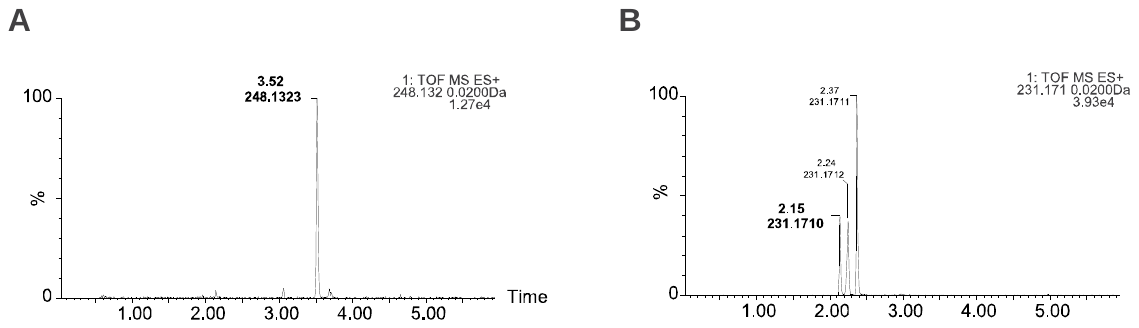

**Supplementary Figure S4.** MS scans of metabolite elutions (A) Zt248 and (B) Zt231, eluting at 3.53 and 2.15 minutes, respectively. The neighboring peaks at the m/z range eluting at different retention times likely represent distinct structural isomers of the same compound.

**Supplementary Tables**

(see separate Excel file)

**Supplementary Table S1:** *Zymoseptoria tritici* field collection information including

collection time point (C1-3), cultivar of origin, field plot.

**Supplementary Table S2:** Relative intensities of 2633 metabolite markers. The isolate name corresponding to sample identifiers are detailed in Supplementary Table S1

**Supplementary Table S3:** List of metabolite-GWAS top significant SNPs and distance to closest gene.

**Supplementary Table S4:** Gene function enrichment based on m-GWAS closest genes to top significant SNPs.
